# A New Tree-Based Methodological Framework to Infer the Evolutionary History of Mesopolyploid Lineages: An Application to the Brassiceae Tribe (Brassicaceae)

**DOI:** 10.1101/2020.01.09.900571

**Authors:** Laura Hénocq, Sophie Gallina, Eric Schmitt, Vincent Castric, Xavier Vekemans, Céline Poux

**Affiliations:** Univ. Lille, CNRS, UMR 8198 – Evo-Eco-Paleo, F-59000 Lille, France

**Keywords:** WGD, mesopolyploids, allopolyploids, phylogenetic inference, homoeologs, orthologs, tree-based orthology inference

## Abstract

Whole genome duplication events are notably widespread in plants and this poses particular challenges for phylogenetic inference in allopolyploid lineages, i.e. lineages that result from the merging of two or more diverged genomes after interspecific hybridization. The nuclear genomes resulting from allopolyploidization contain homologous gene copies from different evolutionary origins called homoeologs, whose orthologs must be sorted out in order to reconstruct the evolutionary history of polyploid clades. In this study, we propose a methodological approach to resolve the phylogeny of allopolyploid clades focusing on mesopolyploid genomes, which experienced some level of genome reshuffling and gene fractionation across their subgenomes. To illustrate our methodological framework we applied it to a clade belonging to the model Brassicaceae plant family, the Brassiceae tribe, that experienced a mesohexaploidy event. The dataset analysed consists of both publically available genomic sequences and new transcriptomic data according to taxa. The present methodology requires a well-annotated reference genome, for which the identification of the parental subgenome fragments has been performed (e.g*. Brassica rapa* and *Brassica oleracea*). Focusing on fully retained genes (i.e. genes for which all homoeologous gene copies inherited from the parental lineages are still present in the reference genome), the method constructs multi-labelled gene trees that allow subsequent assignment of each gene copy to its diploid parental lineage. Once the orthologous copies are identified, genes from the same parental origin are concatenated and tree-building methods are used to reconstruct the species tree. This method allows resolving the phylogenetic relationships (i) among extant species within a mesopolyploid clade, (ii) among the parental lineages of a mesopolyploid lineage, and (iii) between the parental lineages and closely related extant species. We report here the first well-resolved nuclear-based phylogeny of the Brassiceae tribe.

---

Whole genome duplication (WGD) events correspond to large-scale gene duplication processes resulting in the formation of polyploid organisms either by duplication of the whole genome within a given species (autopolyploidy) or by merging of diverged genomes after an interspecific hybridization event (allopolyploidy). WGD events are notably widespread in plants and increasing evidence shows that most extant angiosperm lineages have experienced at least one ancient polyploidization event since the origin of the group (Jiao et al. 2011; Van de Peer et al. 2017). Hence, polyploidy appears to be a fundamental process shaping the evolution and diversification of plant lineages (Otto and Whitton 2000; Marhold and Lihová 2006; Doyle et al. 2008; Soltis et al. 2009; Soltis et al. 2014a; Panchy et al. 2016, Landis et al. 2018).

After an allopolyploidy event, a genome will contain homologous gene copies from different parental origins called homoeologs. Polyploid taxa can be classified as neo-, meso-, or paleo-polyploids according to the age of the last WGD event in their history, and to the degree of subsequent genomic rearrangement (Mandáková et al. 2010). Neopolyploids are the most recently formed polyploids: they display an increase in chromosome numbers and genome size, a highly redundant gene content, and their diploid parents are often still present in the extant flora or fauna. Mesopolyploids have experienced some level of genome reshuffling and gene fractionation (loss of homoeologous copies) across the parental subgenomes, but individual genomic blocks corresponding to each of the parental subgenomes can usually be identified through comparative genomic approaches. The level of gene redundancy in mesopolyploids is generally highly variable among gene ontologies due to differential constraints on functional redundancy and gene dosage flexibility within functional gene networks (Lou et al. 2012; Geiser et al. 2016; Mandáková et al. 2017). The parental nuclear subgenomes of paleopolyploids, the most ancient polyploids, cannot be reliably identified as a result of a strong gene fractionation and profound genomic rearrangements.

Using the nuclear genome for phylogenetic inferences in mesopolyploids is complicated because orthology is difficult to delineate from homoeology. As a result, chloroplast sequences from genic and intergenic regions have been widely used to investigate the phylogenetic relationships within mesopolyploid plant clades (Olmstead et al. 2008, McDill et al. 2009, Warwick et al. 2010; Arias and Pires 2012). The increasing availability of complete chloroplast genomes has further fostered their use for obtaining robust phylogenetic inferences in seed plants (Parks et al. 2009; Guo et al. 2017). However, because the chloroplast genome is usually maternally inherited in flowering plants it will only recover a single parental lineage, which is problematic when applied to allopolyploids. Therefore, the information carried by nuclear genes is not only necessary to detect hybridization and ancient allopolyploidization events but also to recover the whole evolutionary history of polyploid lineages.

Reconstructing the evolutionary history of allopolyploid lineages from nuclear genome sequences requires separating the homoeologous gene copies (i.e. gene duplicates originating from whole genome duplication) into orthologs groups sharing a common origin, i.e. originating from the same ancestral species. Several methods, based on species network reconstruction, have been proposed to reconstruct phylogenies of polyploid lineages with the limitation of being dedicated to neopolyploids, for which at least some of the parental lineages of lower ploidy still exist (Huber et al. 2006; Lott et al. 2009; Albrecht et al. 2012; Jones et al. 2013; Oberprieler 2017; Oxelman et al. 2017). However, none of these methods can be used to reconstruct accurate phylogenies of mesopolyploid lineages for at least three reasons. First, in these lineages, gene copies are often lost differentially among species, which increases drastically the confusion between orthologs and homoeologs. Second, genomes of mesopolyploid species went through a diploidization process and therefore their chromosome counts are not informative. Third, the diploid parental species or lineages are often extinct and therefore cannot be included in the phylogenetic analysis.

Orthology inference methods have been applied to phylogenomic analyses based on transcriptome data. They are often based on reciprocal similarity criterion (e.g. HaMStR, Ebersberger et al. 2009), which could involve wrong homology detection in the case of molecular rate heterogeneity and more problematically in the case of mesopolyploid taxa when a unique copy from different parental origin has remained in different species. Recently, Yang and Smith (2014) proposed a tree-based orthology inference approach capable of accommodating genome duplication in non-model organisms. This method allows reconstructing the phylogeny of mesopolyploid clades, but not to assign homoeologous genes to specific parental lineages. Therefore, the phylogenetic relationships of the mesopolyploid clade with their potential sister groups could not be completely disentangled by this approach.

In this study, we propose a new methodological approach to resolve the phylogeny of allo-mesopolyploid clades. This method requires at least one well-annotated reference genome, member of the studied allo-mesopolyploid clade for which the identification of the parental (progenitors) origin of subgenome fragments has been identified. Focusing on fully retained genes in this reference genome (*i.e.* genes for which one copy is retained in each of the merged parental subgenomes), we construct multi-labelled gene trees (homolog trees) that allow subsequent assignment of each homoeologous gene copy to its diploid parental lineage, *i.e.* separation of orthologs from homoeologs. Once the orthologous copies have been identified within a given subgenome, genes are concatenated and tree-building methods are used to reconstruct the species tree. This method allows us to resolve the phylogenetic relationships *(i)* among all extant investigated species within a mesopolyploid clade, *(ii)* among the parental lineages, which are likely to be extinct lineages and *(iii)* between the parental lineages and closely related extant outgroup species.

In order to test our methodological framework, we applied it to a clade belonging to the Brassicaceae plant family, the Brassiceae tribe. Multiple independent mesopolyploid WGD events have occurred in several Brassicaceae lineages and are likely associated with the observed important species radiation (Mandáková et al. 2017). Broadly, 11 (22%) of the 49 recognized tribes of the Brassicaceae family (Al-Shehbaz 2012) have a mesopolyploid ancestry (Lysak et al. 2005; Lysak et al. 2007; Kagale et al. 2014; Mandáková et al. 2017). In the Brassiceae tribe, the genomes of species analysed to date contain either three or six (in neotetraploid species) copies of orthologous genomic regions of *A. thaliana*. This suggests that the Brassiceae tribe has experienced two successive WGD events, generating a whole genome triplication from which all present-day diploid species in the Brassiceae tribe derive (Lysak et al. 2007). Comparative analyses of the allo-mesohexaploid *Brassica rapa* subgenomes suggested a two-steps origin first involving an allotetraploidization event from two diploid ancestral genomes (named MF1 and MF2, for “Medium fractionated” subgenome and “Most Fractionated” subgenome, respectively), followed by genomic reshuffling and gene fractionation, and then subsequent hybridization with a third diploid parental genome (named LF, for “Least Fractionated” subgenome), again followed by genomic reshuffling and gene fractionation (Cheng et al. 2012; Tang et al. 2012; Cheng et al. 2013; Cheng et al. 2014; Murat et al. 2015). As a result of this complex genomic history, inferring phylogenetic relationships among Brassiceae using nuclear genes is difficult given the high number of homoeologs present in Brassiceae genomes, and differential gene loss/retention following the whole genome triplication. The Brassiceae tribe contains 227 species, 47 genera (Al-Shehbaz 2012) and eight putative monophyletic sub-tribes: “Vella”, “Zilla”, “Cakile”, “Crambe”, “Henophyton”, “Nigra”, “Oleracea” and “Savignya” (Arias and Pires 2012). To date, the reconstruction of phylogenetic relationships within and among clades of the tribe Brassiceae, as well as the tribe circumscription, were performed by using mainly chloroplast markers (Warwick and Black 1991; Warwick and Black 1993; Warwick and Black 1994; Warwick and Black 1997; Arias and Pires 2012). Other markers, such as mitochondrial DNA restriction profiles or nuclear markers (Pradhan et al. 1992; Warwick and Sauder 2005; Warwick et al. 2010; Couvreur et al. 2010; Hall et al. 2011) were more rarely used and reconstructed markedly different topologies. Consequently, a robust Brassiceae phylogeny based on nuclear genes and an appropriate methodology taking into account the mesopolyploid nature of the tribe is still missing.

## Materials & Methods

### Obtaining and Assembling Transcriptomic Data

We used public genomic data for most Brassicaceae outgroup species (*Arabidopsis thaliana*, *Eutrema salsugineum*, *Schrenkiella parvula, Sisymbrium irio*) and for all Brassiceae available (*Raphanus raphanistrum, Raphanus sativus, Brassica nigra, Brassica rapa, Brassica oleracea*) (Table 1). The well-annotated genomes of *B. rapa* and *B. oleracea* were used as reference genomes for the analyses.

**Table 1.**
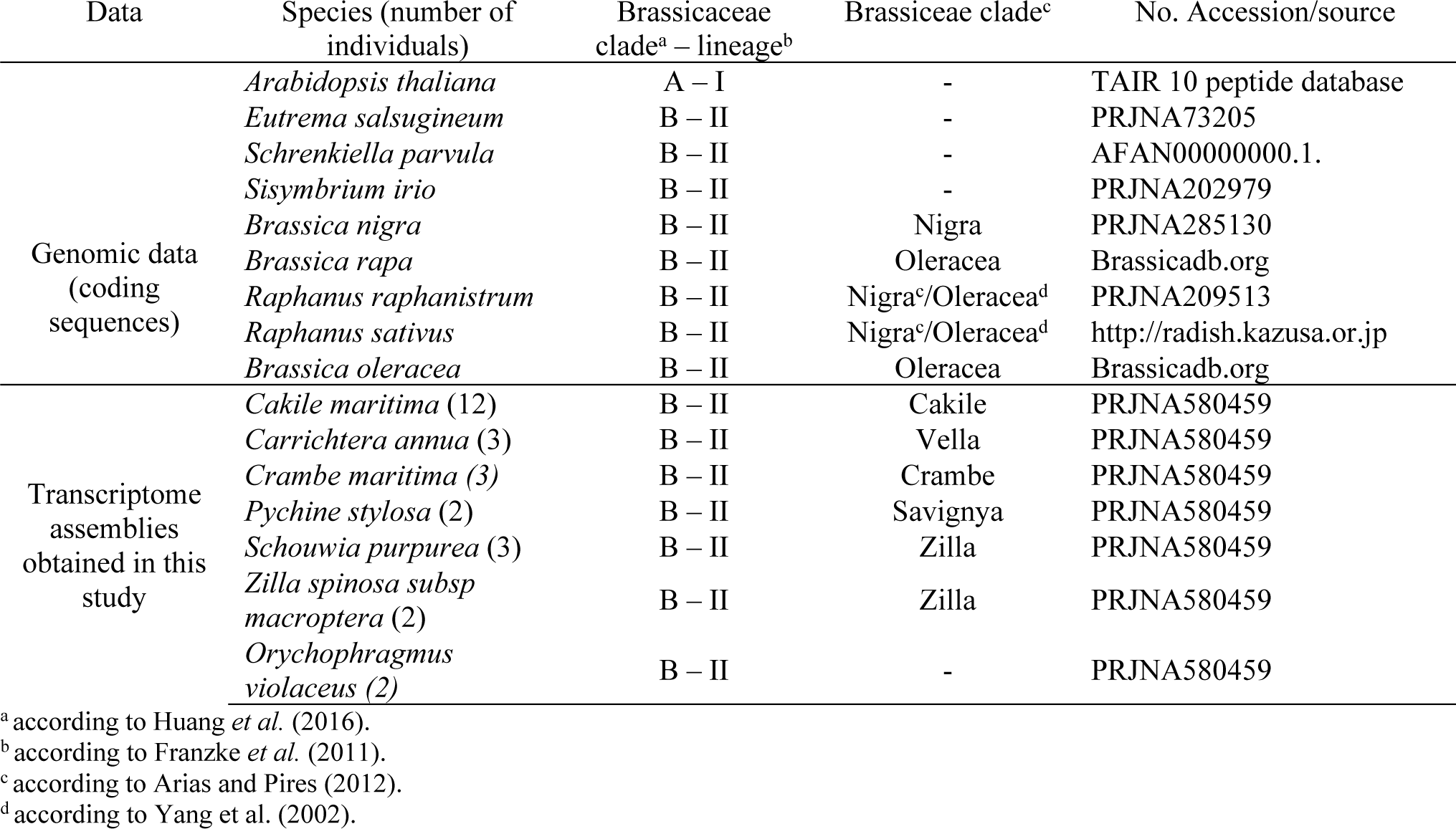
Genomic data and transcriptome assemblies used for the present study. Numbers between brackets represent the number of individuals sequenced for a given species.

#### Plant material

In addition to genomic data, we obtained transcriptomic sequence data for the Brassicaceae outgroup species *Orychophragmus violaceus* (L.) O.E. Schulz, and 6 Brassiceae species representative of the various sub-tribes: *Carrichtera annua* (L.) DC. (clade Vella), *Zilla spinosa subsp. macroptera* (Coss.) Maire & Weiller (clade Zilla), *Schouwia purpurea* (Forssk.) Schweinf. (clade Zilla), *Psychine stylosa* Desf. (clade Savignya), *Cakile maritima* Scop. (clade Cakile) and *Crambe maritima* L. (clade Crambe). Flower buds of *Cakile maritima* and *Crambe maritima* were collected in natural populations at the Digue du Braek (Dunkerque, France) and at the mouth of the Slack river (Ambleteuse, France), respectively. Flower buds of *O. violoceus* were collected at the National Botanic Garden of Belgium (Meise). Seeds of all other species were obtained from the Plant Gene Resources of Canada and originated from various botanical gardens (online Appendix 1, available from the Dryad Digital Repository at http://dx.doi.org/10.5061/dryad/[NNN]). Seeds from each species were sown in potting soil and germinated at 20°C in a greenhouse providing a 12-h photoperiod for several weeks and controlled conditions until germination. Plants were grown in greenhouse conditions until flowering. Flower buds were collected for RNA sequencing, except for the two individuals of *Zilla spinosa subsp. macroptera* for which the RNA extraction was performed from leaves due to the lack of flowering. We sequenced 2 to 12 individuals per species (online Appendix 1, available on Dryad) according to availability.

#### cDNA library preparation and transcriptome sequencing

Total RNA was extracted from flower buds or leaves with the Spectrum Plant Total RNA kit (Sigma, Inc., USA), following the manufacturer’s protocol, and treated with DNAse (On-Column DNase I Digestion set, Sigma, Inc., USA). cDNA libraries were prepared with the TruSeq RNA sample Preparation v2 kit (Illumina Inc., USA). Each cDNA library was sequenced using a paired-end protocol on HiSeq2000, HiSeq2500 or HiSeq3000 sequencer, producing 100 to 150-bp reads (twelve libraries pooled in equi-proportion per lane) (online Appendix 1, available on Dryad). Raw reads were submitted to the SRA database under the accession number PRJNA580459. Demultiplexing was performed using CASAVA 1.8.1 (Illumina Inc., USA) to produce paired sequences files containing reads for each sample in the Illumina FASTQ format. RNA extraction, library preparation, and sequencing were done by the sequencing platform in the AGAP laboratory, Montpellier, France (http://umr-agap.cirad.fr/).

We then used FastQC (Andrews 2010), a quality control tool for high throughput sequence data. Adaptor sequences and poly-A tails were trimmed and reads showing GC content bias, low complexity, small size or exact duplicates were removed using PRINSEQ (Schmieder and Edwards 2011) and CUTADAPT (Martin 2011). We controlled the quality of cleaned reads by using FastQC again (online Appendix 1, available on Dryad).

#### De novo transcriptome assembly

Transcriptome assembly was performed using TRINITY with default parameters (Grabherr et al. 2011). To minimize redundancy in each transcriptome assembly (mainly due to isoforms), we used CAP3 to generate consensus sequences (Huang and Madan 1999). Overlaps of 120bp and 98% of identity between two or more contigs induced the construction of consensus sequences. Then, we used QUAST (Gurevich et al. 2013) to evaluate the assembly quality (online Appendix 2, available on Dryad).

### Assigning Nuclear Homoeologous Gene Copies to Parental Subgenomes

An overview of the developed method is displayed in Figure 1. In the main text analyses are described with more details.

**Figure 1.**
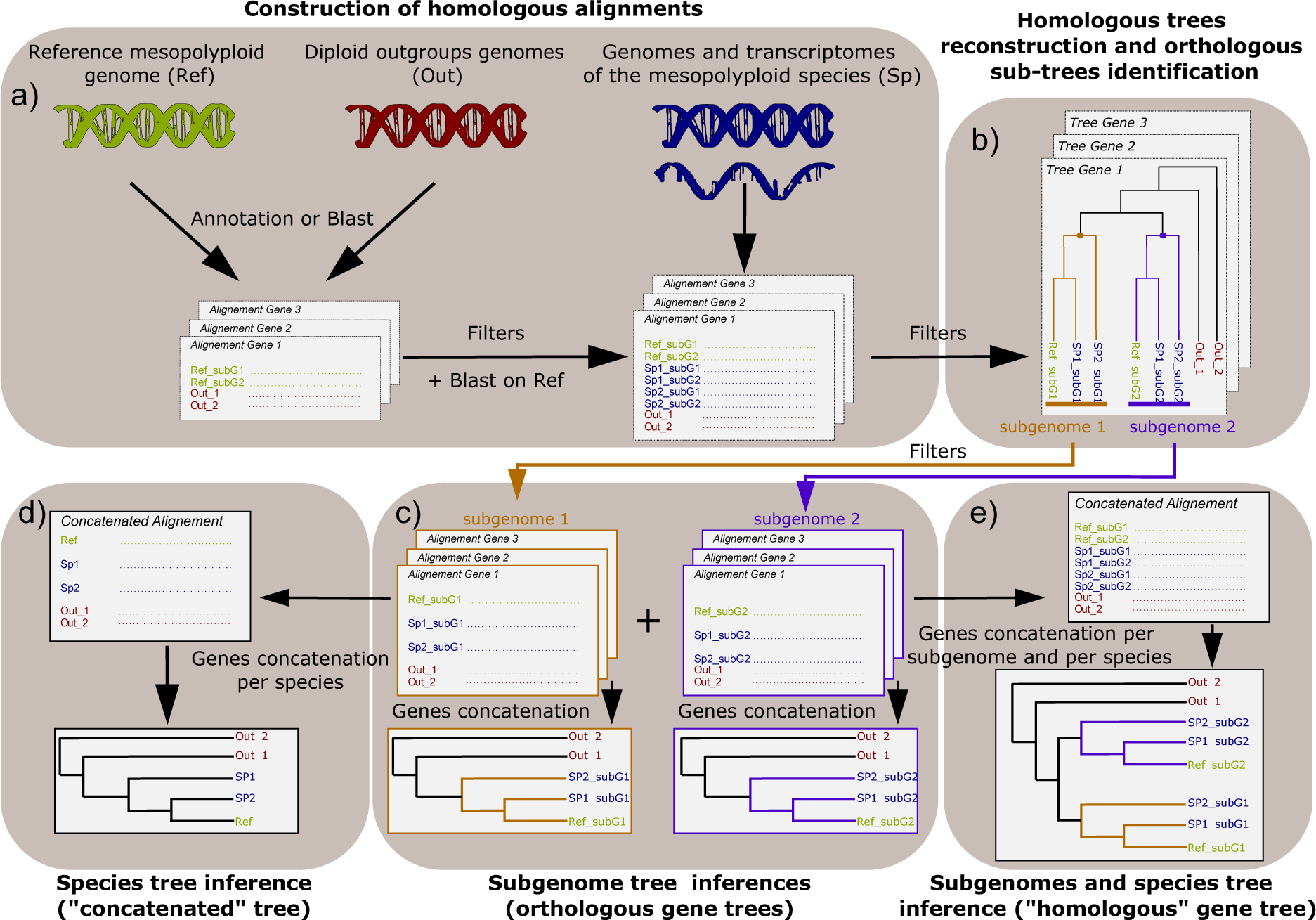
Methodological framework developed in this study for reconstructing the evolutionary history of mesopolyploid lineages. For the sake of simplicity, in this sketch a mesotetraploid plant example was used. Several phylogenetic trees are obtained: a tree for each subgenome based on orthologous gene copies (c), a species tree (d) and a homologous tree based on both orthologous and homoeologous gene copies (e). “subG” refers to subgenome.

#### Subgenome annotation database

We focused on the specific genes retained as three homoeologous copies (referred to here as “triplets”) in both *B. rapa* and *B. oleracea* reference genomes, representing the so-called LF (least-fractionated), MF1 (medium-fractionated) and MF2 (most-fractionated) subgenomes. This represented a starting set of 1,344 genes, referred to here as the Liu database (LiuDB) (Liu et al. 2014). We retrieved *Arabidopsis thaliana* orthologous sequences from the TAIR database and excluded genes whose *A. thaliana* ortholog was shorter than 500 bp, leading to 1,163 remaining genes. We then retained only *B. rapa* and *B. oleracea* gene sequences longer than 500bp and with a length coverage of the orthologous *A. thaliana* sequence higher than 60%. This step resulted in a final set of 1,085 homologous genes, each containing the three homoeologous copies of *B. rapa* and of *B. oleracea* as well as the single orthologous copy of *A. thaliana* (Fig. 1, panel 1). The term homologous is used when homoeologous and orthologous gene copies are both present in the same dataset or phylogenetic tree.

#### Orthologous gene sorting

For each set of homologs, triplets of sequences of *B. rapa* and *B. oleracea* and the sequence from *A. thaliana* were aligned with MACSE (Ranwez et al. 2011). Alignments were trimmed on both sides following the *A. thaliana* sequence and poorly aligned sites were removed using trimAl v1.2 with default settings (Capella-Gutiérrez et al. 2009). When the final alignment was at least 500bp long, phylogenetic trees were built with RAxML (Stamatakis 2014) with a GTR+Γ model of sequence evolution. We discarded all homolog trees in which the *B. rapa* and *B. oleracea* copies originating from the same subgenome (LF, MF1 or MF2) were not monophyletic – which may arise due to annotation errors or ectopic gene conversion events between homoeologs (Soltis et al. 2014b; Scienski and Conant 2015). In addition, phylogenetic trees missing one or more subgenomes, due to sequence elimination during alignment trimming, were discarded. For each of the 1,075 remaining alignments of homolog gene copies, single orthologous sequences from the three additional outgroup species and up to three homoeologous sequences from each Brassiceae were added to the alignments as explained below. Each Brassiceae species in the final alignments was represented by 1 to 12 individuals (Table 1).

For the three additional Brassicaceae outgroup species, *Eutrema salsugineum* (Yang et al. 2013), *Schrenkiella parvula* (Dassanayake et al. 2011) and *Sisymbrium irio* (Haudry et al. 2013), we extracted orthologous sequences to the 1,075 selected *A. thaliana* gene sequences using BLASTN (Altschul et al. 1990) (minimum percentage of identity: 80%; minimum length: 60% of the reference sequence). For the Brassiceae species and *O. violaceus* (phylogenetically close to the Brassiceae and potentially sharing a WGD event with them [Lysak et al. 2007]), the homoeologous genes present in three copies in *B. rapa* and *B. oleracea* were mapped using BLASTN (minimum percentage of identity: 80%; minimum length: 60% of the reference sequence) onto each of our original transcriptome assemblies and onto the genome sequences of three additional Brassiceae species: *Raphanus sativus* (Kitashiba et al. 2014), *Raphanus raphanistrum* (Moghe et al. 2014) and *Brassica nigra* (Yang et al. 2016) (Fig. 1a). In order to avoid the introduction of chimeric contigs – particularly from transcriptomes – we only extracted the portion of each best-hit contig that aligned with the reference sequence. Because of alternative splicing, several isoforms can be the best-match sequence of the same Brassica reference sequence. We only extracted the longest isoform. The alignment and trimming were done as previously explained with the “Arabidopsis - Brassica” homologous datasets and led to 1,044 filtered alignments of homolog gene sequences. Table 1 summarizes the data type and their source as well as the species included in the present study.

For each of the 1,044 filtered alignments of homolog gene sequences, we identified each of the three expected sub-trees of orthologs by performing node annotations (Fig. 1b) with the ETE v3 toolkit (Huerta-Cepas et al. 2016). First, the two reference sequences of a given subgenome were localized on the tree (*e.g.* for the subgenome MF1, we localized the *B. rapa* and *B. oleracea* MF1 sequences). Then, from the node representing the common ancestor of these two reference sequences, we climbed backward through the tree until reaching the last node defining a clade containing neither outgroup sequences (including *O. violaceus*) nor reference sequences from other subgenomes (*e.g.* in this case MF2 or LF sequences). Finally, we annotated the corresponding node with the subgenome label of the *B. rapa* / *B. oleracea* reference sequences. The process was applied on each homolog tree in order to annotate the three sub-trees corresponding to the three subgenomes (LF, MF1 and MF2). Homolog trees with more than 10% of Brassiceae transcriptome sequences localized outside the defined orthologs’ sub-trees were discarded. A total of 867 remaining homolog trees was obtained.

### Species Tree Inference using Nuclear Genes

Once the orthologs’ sub-trees were defined, corresponding sequences were extracted from the alignments according to some filters (see below), written in three distinct files representing each orthologs’ group and aligned (Fig. 1c). For a given species, only the longest sequence was extracted under the condition that all sequences of this species were monophyletic in a given orthologs’ sub-tree. If not, no sequence was extracted for this species in the concerned orthologs’ sub-tree (232 cases). Moreover, orthologs’ sub-trees in which there were less than two Brassiceae transcriptome sequences belonging to two different species were filtered out (140 sub-trees discarded). The tree topologies indicated that the mesotetraploid *O. violaceus* (Lysak et al. 2007) did not share any WGD event with the Brassiceae clade. For this species, we therefore extracted only one sequence (the longest) from each homolog dataset.

Orthologs’ alignments were concatenated in different ways: first, separately in a LF matrix (839 genes; 1,000,401 bp), a MF1 matrix (829 genes; 982,048 bp) and a MF2 matrix (814 genes; 962,921 bp) (Fig. 1c); second, all together in a “concatenated” matrix (2,482 genes; 2,945,370 bp) (Fig. 1d). Finally, in order to resolve the evolutionary history of the mesopolyploid group, we built a fifth matrix, called “homologous”, in which all the homoeologous gene copies of a given Brassiceae species were represented as a multi-labelled tree with subgenome labels (757 genes, 881,473 bp) (Fig. 1e).

For each DNA alignment, the best partitioning scheme was assessed by PartitionFinder 2 with the rcluster search mode using the corrected Akaike Information Criterion (AICc) according to Lanfear et al. (2017). A phylogenetic species tree was then constructed for each of the five matrices (Fig. 1c to 1e) according to the best-partition scheme selected by PartitionFinder 2. Maximum Likelihood (ML) analyses conducted with RAxML v8.2.10 (Stamatakis 2014) were run under a GTR+Γ model of sequence evolution for each DNA partition. The number of bootstrap replicates was automatically determined (Stamatakis et al. 2008). Bayesian inference (BI) analyses were performed using MrBayes v3.2.6 (Ronquist et al. 2012) with a different model for each DNA partition. Two runs of 75,000,000 generations were completed with four chains each and trees were sampled every 2,000 generations for the LF, MF1, MF2, and “homologous” matrices. 165,000,000 generations were completed for the “concatenated” alignment. Plots of the likelihood-by-generation were drawn to check chain convergence, indicated as well by an average standard deviation of split frequencies smaller than 0.01, a Potential Scale Reduction Factor at 1.0 and effective sample size values above 100. The first 25% of trees from all runs were discarded as burn-in. A majority-rule consensus of the remaining trees from both runs was used to obtain the posterior probability tree.

### Influence of missing data

The proportion of alignment gaps for each species and each matrix (LF, MF1, MF2 and “concatenated”) is reported in online Appendix 3 (available on Dryad) and the number of gene sequences for each Brassiceae species is indicated in online Appendix 4 (available on Dryad). Alignment gaps can be due to indels, gene copy deletion or missing data (due in part to low expression level of a given homoeologous gene copy). Therefore, the occurrence of a high proportion of alignment gaps indicates a high proportion of missing information, which could impair the accuracy of the phylogenetic inference (Lemmon et al. 2009). To test the influence of the missing data, we produced four additional filtered matrices (LF, MF1, MF2 and “concatenated” filtered-matrix) in which sites displaying missing data for at least one of the four following species *C. annua*, *P. stylosa*, *Z. spinosa subsp. macroptera* and *S. purpurea* were removed. We focused on these four species because they displayed a large amount of missing data (45 to 71% of alignment gaps, depending on the matrix, online Appendix 3, available on Dryad). The lengths of the resulting matrices were 183,904; 77,944; 72,187 and 334,035 bp for the LF, MF1, MF2 and the “concatenated” filtered-matrices, respectively. Phylogenetic analyses were conducted as previously mentioned. For the Bayesian inferences, 10,000,000 (LF, MF1, MF2 filtered-matrices) to 30,000,000 (“concatenated” filtered-matrix) generations were completed and trees were sampled every 400 generations.

### Influence of the Subgenome Annotation Database

Our method relies on the subgenomes annotation of the mesopolyploid species used as a reference (here, *B. rapa* and *B.oleracea*). Most of the published studies use a synteny approach for assigning chromosomal fragments to their parental genome. However, there are some substantial differences among studies in the way homoeologs are assigned to a given subgenome, and this results in variation in the number of genes with homoeologous copies for each of the three subgenomes in *B. rapa*: 1,578 in Wang et al. (2011a) (genome used in the analysis of Liu et al. 2014), 1,675 in Cheng et al. (2012) and 506 in Murat et al. (2015) as well as in the identity of these genes. We thus investigated to which extent the database of Liu et al. (2014) (LiuDB) used in our study would give concordant results with the database of Murat et al. (2015) (MuratDB). To that end, we compared results when our new methodology was applied either to the entire Murat database or to a restricted dataset containing only the concordant triplets between the Liu and the Murat databases, i.e. triplets sharing the same annotation in both databases (the “Concordant Triplets” dataset - CTDB, see online Appendix 5 for a detailed protocol, available on Dryad).

To evaluate statistical support for incongruent phylogenetic topologies, we used the program SOWHAT (Church et al. 2015) that implements the Swofford-Olsen-Waddell-Hillis (SOWH) test (Goldman et al. 2000). The test was performed following the recommendation of the authors, with a number of sample replicates of 1,000, using RAxML and the model GTR+Γ for both inference and simulation. Data were simulated with the same number and position of gaps compared to the original data set and under a zero-constrained tree (Susko 2014). The most likely topology inferred from the “homologous” alignment obtained using either the Murat or the “Concordant Triplets” databases was tested against the alternative topology obtained with the Liu database. The reverse tests were performed as well.

## Results

### Transcriptome Assemblies

The transcriptome of 26 individuals from six different species of Brassiceae and two individuals of the outgroup *Orychophragmus violaceus* were sequenced (Table 1) and paired-end Illumina reads were assembled after cleaning. According to the species and the tissue sampled, the transcriptome assemblies yielded between 28,710 and 57,725 contigs (online Appendix 2, available on Dryad). The total contigs length for a given transcript varies between 38,803,080 and 80,126,538 bp, the N50 varies between 1,596 and 1,857 bp and the percentage in GC varies between 42.11 and 43.84 % (online Appendix 2, available on Dryad). Both *Zilla spinosa subsp macroptera* individuals, for which leaf tissues instead of flower buds were used, displayed the lowest total sequence length and the lowest number of contigs. This was expected as reproductive organs have a much broader range of expression than leaves (Schmid et al. 2005).

### Gene Orthology Assignment

Starting with the 1,085 genes longer than 500 bp that were present in three copies in *B. oleracea* and *B. rapa* and with a sequence coverage higher than 60% with the orthologous sequence of *A. thaliana*, we obtained a dataset of 867 genes for which each homoeologous copy sequenced in a Brassiceae species could be assigned to a specific parental subgenome. The remaining 218 genes were removed after applying the filters described in the methods section.

### Phylogenetic Reconstruction of a Mesopolyploid Clade using Nuclear Coding Sequences

The methodology developed here allowed us to reconstruct a phylogenetic tree for each of the parental subgenomes, but also to place them on the same phylogenetic tree, thus revealing their evolutionary history (Fig. 1).

The intra-Brassiceae phylogenetic relationships are fully congruent and robust, with high support values at each node, whatever the dataset (individual or concatenated subgenomes) or the phylogenetic reconstruction method used (Fig. 2 and online Appendix 6). “Vella” is the first diverging clade followed by “Zilla”. The “Savignya” sub-tribe appears to be the sister group of the core Brassiceae (“Oleracea” + “Nigra” + “Crambe” + “Cakile”). The “Crambe”, “Nigra” and “Oleracea” clades share a common ancestor and the latter two are most closely related to each other.

**Figure 2.**
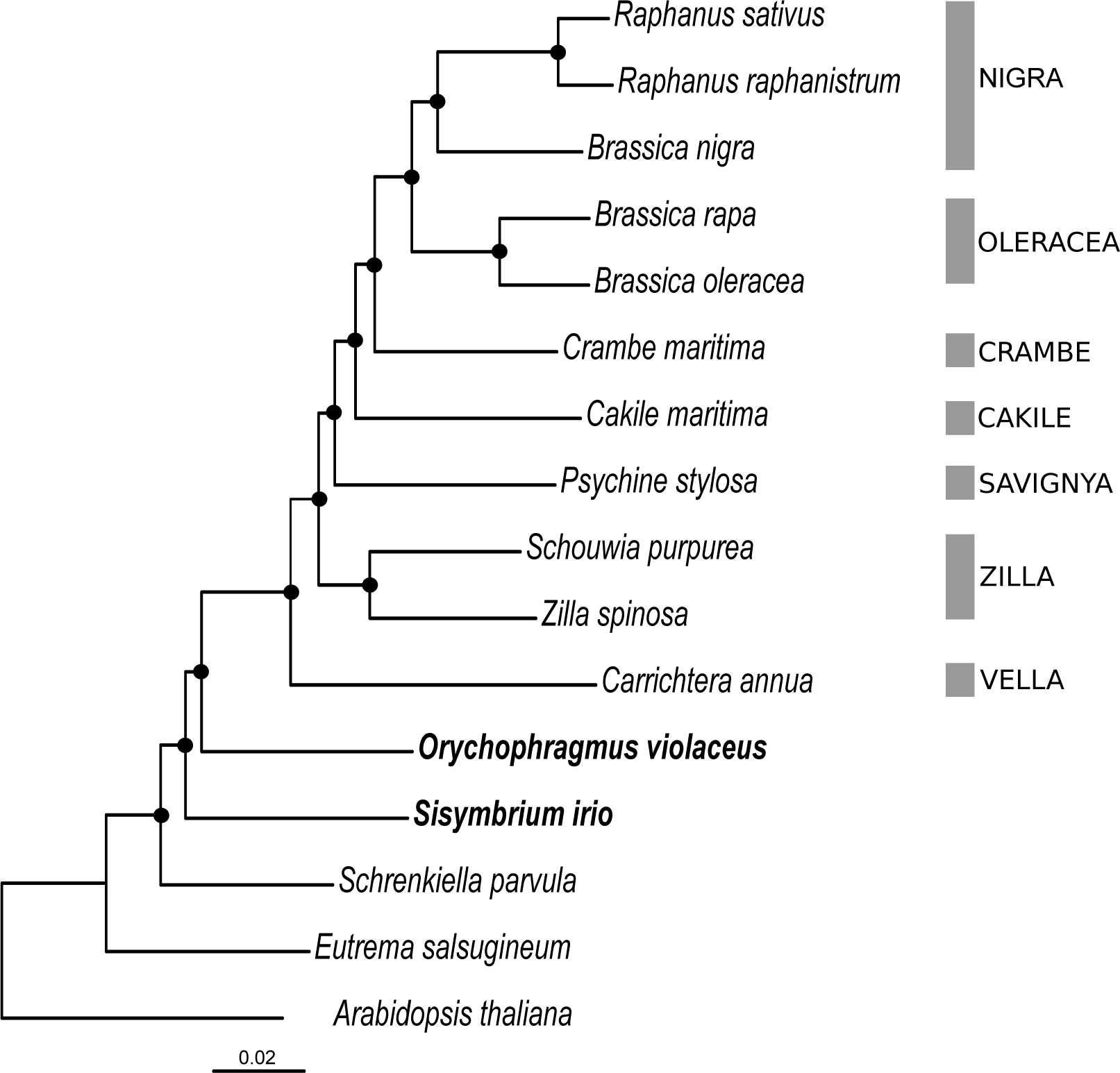
Maximum likelihood phylogeny of the tribe Brassiceae based on the “concatenated” nuclear genes alignment (concatenation of the LF, MF1 and MF2 matrices). The seven investigated Brassiceae sub-clades are indicated on the right. Brassiceae’ closest outgroups are in bold. Black circles indicate maximal support values for both ML and Bayesian analyses (BP = 100 / PP = 1.0).

The phylogenetic positions of *O. violaceus* and *S. irio* were more uncertain and varied according to the dataset used: each subgenome displayed a different topology (Fig. 3). The position of these two species in the “concatenated” species tree was similar to that of the MF1 tree (Figs. 2 and 3).

**Figure 3.**
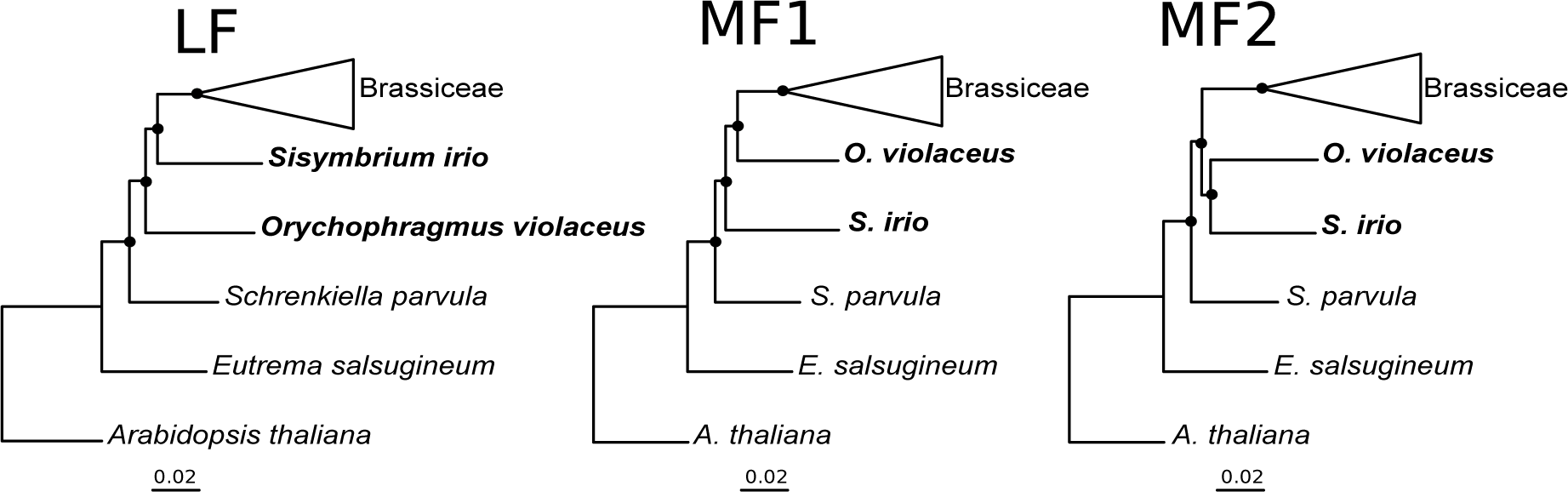
Maximum likelihood phylogenies of the Brassiceae and their relatives based on the LF, MF1 and MF2 subgenomes data. Brassiceae’ closest outgroups are in bold. Black circles indicate nodes with maximal support (BP = 100 / PP = 1.0).

Trees obtained from the filtered matrices, to assess the influence of missing data, yielded largely identical topologies, although nodes support values dropped compared to the full-matrix trees (online Appendix 7, available on Dryad). The only exception was the relative position of the outgroup species *S. irio* and *O. violaceus*, exchanged in the “concatenated” species tree after filtering (Fig. 2 and online Appendix 7, available on Dryad).

The species tree obtained with the “homologous” alignment (836,958 bp) displaying all three lineages LF, MF1 and MF2 is reported in Fig. 4. The fully resolved topology of the Brassiceae species in each of the three sub-trees was congruent with all other inferred topologies (“concatenated”, LF, MF1 and MF2) (Fig. 2 and online Appendix 6). Interestingly, *O. violaceus* and *S. irio* appear to be more closely related to the LF parental subgenome than to the other two subgenomes, and their relative position is congruent with the LF tree (Fig. 3). In the same way, the MF2 subgenome diverges equally from *S. irio* and *O. violaceus*, which is congruent with the topology of the MF2 tree (Fig. 3). However, the MF1 subgenome appears to be sister group to the MF2 clade and therefore equally distant from *S. irio* than from *O. violaceus,* which is not congruent with any of the MF1-filtered or -unfiltered trees (Fig.3 and online Appendix 7, available on Dryad).

**Figure 4.**
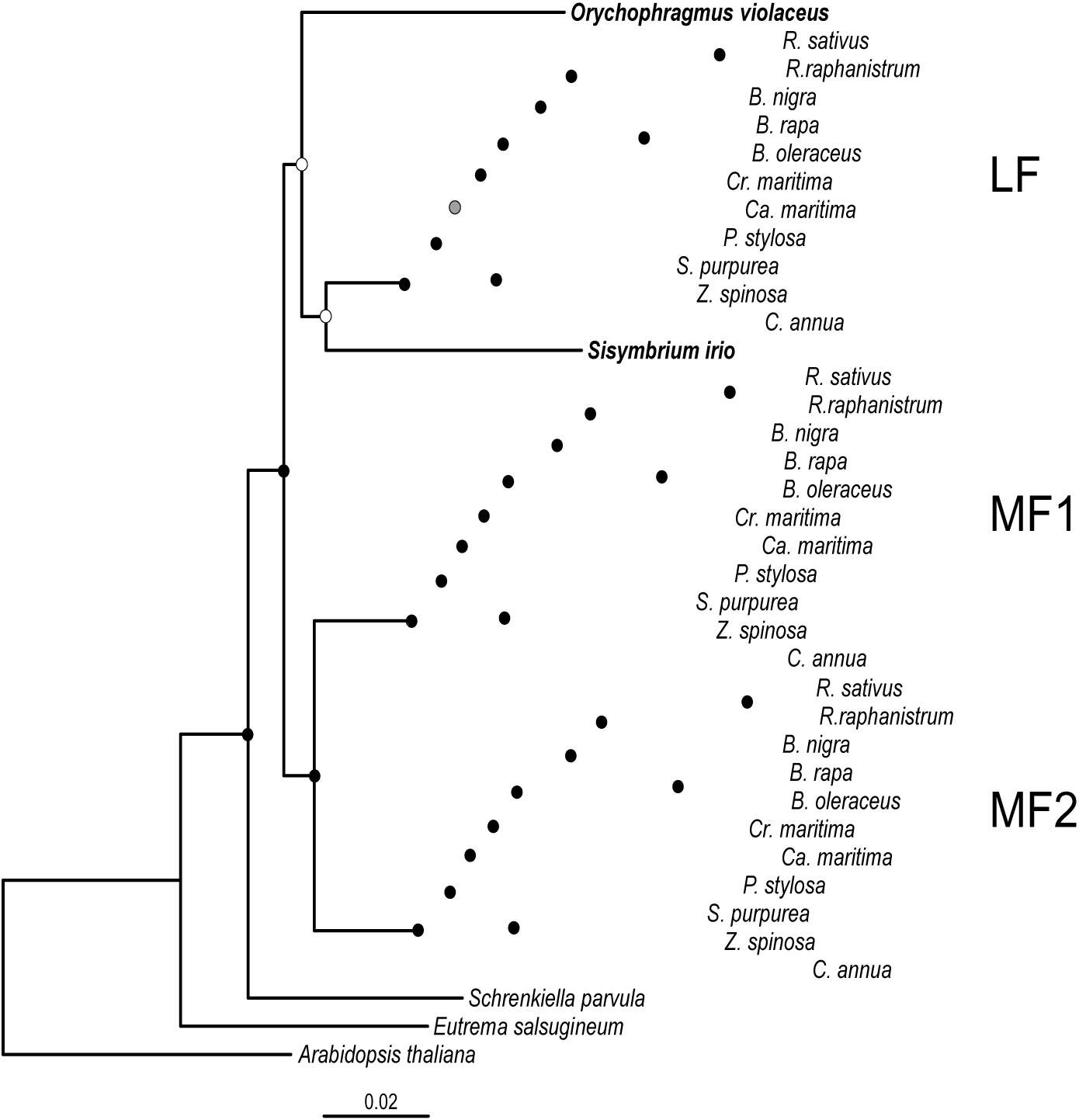
Maximum likelihood phylogeny of the tribe Brassiceae based on the “homologous” alignment obtained with the Brassica genes annotation of Liu et al. (2014). The phylogeny displays the three sub-trees corresponding to the three subgenomes present in the Brassiceae species. Brassiceae’ closest outgroups are in bold. Black circles indicate nodes with maximal support (BP = 100 / PP = 1.0). Smaller support values are indicated with grey (90 ≤ BP <100 and PP=1.0) and white (80 ≤ BP < 90 and PP=1.0) circles.

### Influence of the Subgenome Annotation Database

In order to assess the influence of the subgenome annotation database on the results, we applied our pipeline on the *B. rapa* data from Murat et al. (2015) (MuratDB). We obtained 472 gene triplets. From these triplets, 158 (33.5%) were absent from the LiuDB, either because we considered only triplets present in both *B. rapa* and *B. oleracea* or because some genes found as triplicates in the MuratDB were found as duplicates or single copy in Liu et al. (2014). Only 181 of the 472 triplets (38.3%) were present in both datasets with exactly the same annotation for each of the three subgenomes (“concordant triplets”). For a large part, the discordance found for the 133 remaining triplets (28.2%) was either due to inversions in the subgenomes annotation between LF, MF1 and MF2 (∼4/5) or to differences in the identity of homoeologs between triplets (∼1/5).

We reconstructed the phylogeny of the tribe Brassiceae with both the MuratDB (196 / 472 remaining genes after all filtering steps) and the CTDB datasets (120 / 181 remaining genes) (online Appendix 5, available on Dryad). The total size and the number of genes of each concatenated alignment are reported in Table 2. The topologies obtained were similar to those using the LiuDB, although with generally lower support for some branches (Fig. 5 and online Appendix 8 to 10, available on Dryad). One notable exception was the phylogenetic relationships between the three Brassiceae subgenomes and their close relatives on the “homologous” trees: *S. irio* and *O. violaceus* were external groups of all three parental lineages with both the MuratDB and the CTDB datasets (Fig. 5 and online Appendix 10, available on Dryad) whereas they were external groups of the LF subgenome only with the LiuDB (Fig. 4). As a consequence, both “homologous” trees were only congruent with the corresponding LF topology but not with the MF1 nor with the MF2 ones (Fig. 5 and Online Appendix 8 to 10, available on Dryad).

**Figure 5.**
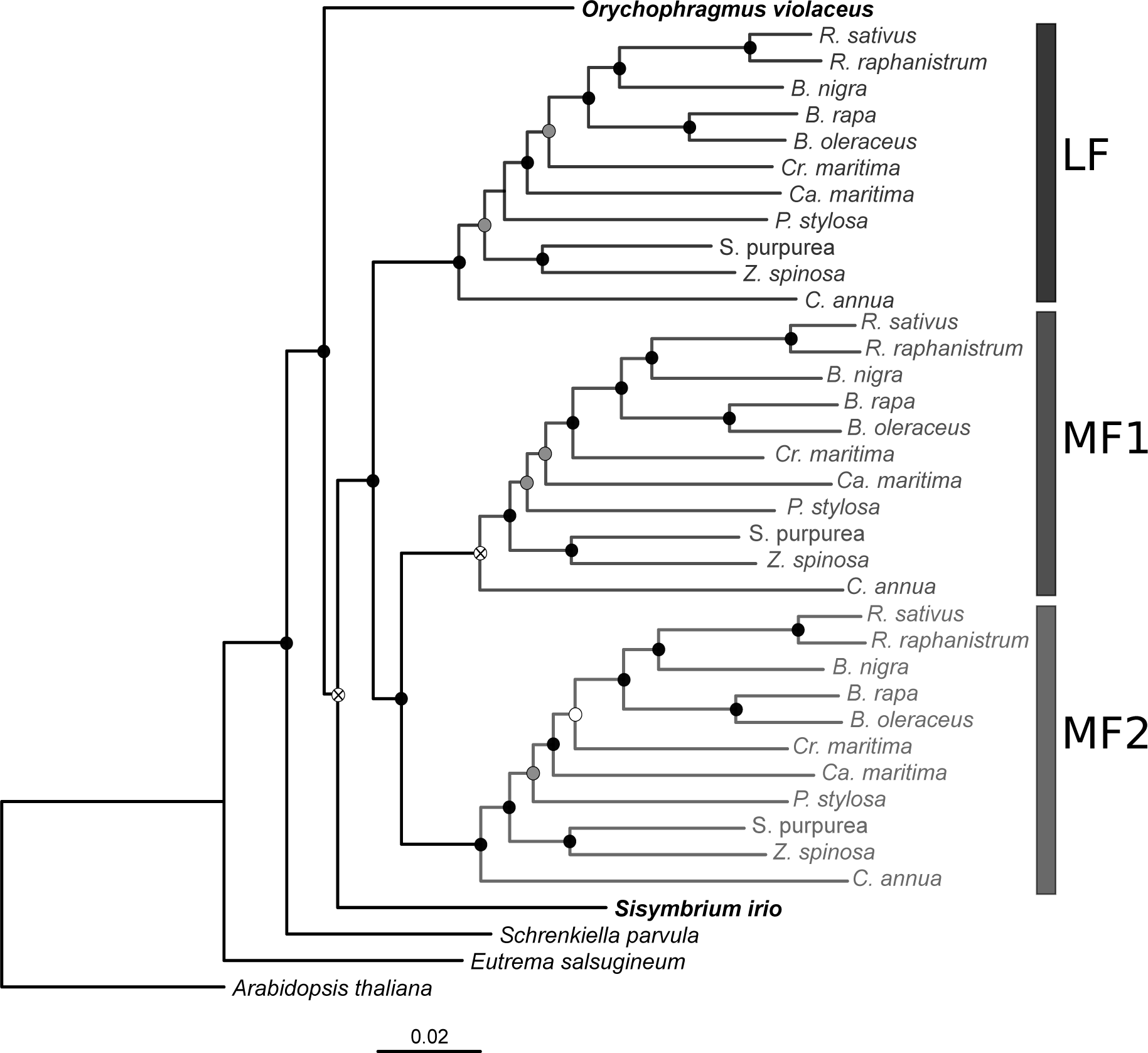
Maximum likelihood phylogeny of the tribe Brassiceae based on the “homologous” alignment obtained with the Brassica genes annotation of Murat et al. (2015). The phylogeny displays the three sub-trees corresponding to the three subgenomes present in the Brassiceae species. Brassiceae’ closest outgroups are in bold. Black circles indicate nodes with maximal support (BP = 100 / PP = 1.0). Smaller support values are indicated with grey (90 ≤ BP <100 and PP=1) and white (80 ≤ BP < 90 and PP=1.0) circles. Supports with a BP < 80 and a PP=1.0 are indicated with a cross.

**Table 2.** The size (in base pairs) and the number of genes for each alignment inferred from the genome annotation of Murat et al. (2015) (MuratDB) and from the “concordant triplets”, i.e. triplets with the same subgenome annotation in Liu et al. (2014) and in Murat et al. (2015) (CTDB).

| Matrix | MuratDB | CTDB |
| --- | --- | --- |
| LF | 220,428 bp, 185 genes | 143,076 bp, 115 genes |
| MF1 | 225,237 bp, 191 genes | 148,219 bp, 119 genes |
| MF2 | 218,648 bp, 186 genes | 136,014 bp, 110 genes |
| “concatenated” | 664,313 bp, 562 genes | 427,309 bp, 344 genes |
| “homologous” | 198,705 bp, 172 genes | 127,939 bp, 104 genes |

SOWH tests were used to compare the alternative phylogenetic relationships between the two closest outgroups (*S. irio* and *O. violaceus*) and the three parental lineages among the three databases (LiuDB, MuratDB and CTDB). In all cases, the outcome of the SOWH tests were similar and indicated that the best topology obtained with a given dataset significantly overpassed the tested alternative topologies. The minimum confidence intervals fall entirely on one side of the significance level of 0.01, suggesting that the observed discrepancy is not due to insufficient phylogenetic signal or small sample sizes (Table 3).

**Table 3.**
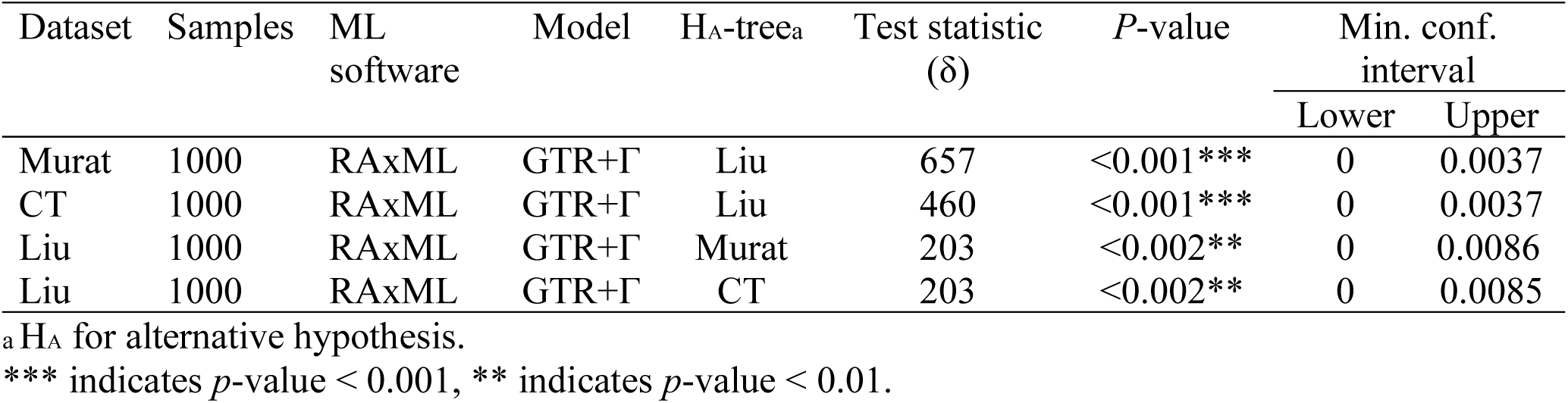
Comparison of alternative phylogenetic hypotheses. SOWH tests were used to determine whether the ML tree from Murat and CT datasets differs significantly from the ML tree inferred from Liu dataset (constrained tree) (first two lines). Conversely, the test was also performed in order to determine whether the ML tree from the Liu dataset differs significantly from the ML tree inferred from Murat and CT datasets (constrained tree) (last two lines).

## Discussion

### Using Fully Retained Genes for Phylogenetic Analyses within Mesopolyploid Lineages

Allopolyploid lineages are formed by the merging of divergent genomes, and as such they represent a particular challenge for phylogenetic analysis because homoeology can be hard to distinguish from proper orthology. Here, by focusing on genes for which all homeologs have been retained, we aimed at assigning gene copies of polyploid lineages to their respective parental subgenomes in order to build orthologous groups that we can subject to proper phylogenetic analyses. Moreover, performing separate phylogenetic analyses for each parental subgenome allows testing for congruence among the inferred topologies, which brings insight into the quality of the obtained topology. Application of our method to the mesohexaploid Brassiceae tribe was very successful as we obtained strictly consistent topologies of the Brassiceae sub-clades among the three parental subgenome analyses, and these results were robust to missing data or the choice of subgenome annotation method. This approach is particularly well suited for mesopolyploid lineages, i.e. polyploid lineages that have experienced some level of genome reshuffling and gene fractionation across the parental subgenomes, but for which individual genomic blocks corresponding to each of the parental subgenomes can still be identified. Indeed, in such groups, gene copies are not necessarily orthologs and ignoring this fact would generate strong noise in phylogenetic reconstruction. For example, in the mesohexaploids *B. rapa* and *B. oleracea*, 46.3% of ancestral genes are retained in multiple copies with 10.5% of fully retained genes (3 copies) (Liu et al. 2014). Similarly, in the mesotetraploid maize genome, 18% of ancestral genes are retained as duplicates (Schnable et al. 2011).

The strength of our approach would be maximized if the same set of ancestral genes was fully retained in all clades that share the mesopolyploidy event, allowing to generate full sets of orthologous alignments and minimizing missing data. Convergent patterns of gene loss versus gene retention have been commonly reported in mesopolyploid genomes (Barker et al. 2008; Haudry et al. 2013; Li et al. 2016; Moghe et al. 2014; Geiser et al. 2016; Mandáková et al. 2017). These convergent patterns of retention of functional groups of genes are thought to be caused mainly by gene dosage balance constraints (Birchler and Veitia 2007; Freeling 2009). However, in our dataset, many copies of the Brassica homoeologous triplets were missing in the other investigated Brassiceae species, regardless of the type of data (genomic or transcriptomic, Supplementary Table S3, available on Dryad). The absence of one or more copies in a given species can be explained by lineage-specific losses during the gene fractionation process, the lack or low level of expression (for transcriptomic data) as differential expression among homoeologous copies is common (Cheng et al. 2012) and/or incorrect assembly of raw sequencing reads. Lineage-specific gene conversion events between homoeologs could also explain that we failed to recover some of the Brassica homoeologous copies used as references (Wang et al. 2009; Wang et al. 2011b; Wang and Paterson 2011; Scienski and Conant 2015). However, our analyses showed that the obtained topologies do not seem to be impacted by the amount of missing data.

An important issue is the quality of the annotation of the separate parental subgenomes used as references. Although the identification of homoeologous blocks in well-assembled mesopolyploid genomes is rather straightforward, the assignment of each block to its respective parental subgenome may be error-prone. Such annotation generally assumes different levels of fractionation (biased gene loss) among subgenomes, with the implicit assumption that the level of fractionation is homogenous within each subgenome (Wang et al. 2011a). Accordingly, our approach will be best suited to cases with strong biased gene fractionation, because this will decrease the possibility of erroneous annotations. Analysis of 15 genomes descending from six ancient eukaryotic WGD events suggested that biased gene fractionation is a universal feature across eukaryotes (Sankoff et al. 2010), and could be caused by differential overall level of gene expression among subgenomes, a phenomenon known as genome dominance (Schnable et al. 2011). Biased gene fractionation seems also to be common in flowering plants (e.g. Arabidopsis, Thomas et al. 2006; maize, Schnable et al. 2011; Brassica, Wang et al. 2011a; cotton, Zhang et al. 2015).

### Reconstructing the Evolutionary History of Diploid Parental Lineages

Another asset of our approach is that it can reveal the evolutionary history of the diploid parental lineages which cannot be analyzed simultaneously because they are usually extinct. In our study, using the Liu et al. (2014) annotation, we found that the MF1 and MF2 parental lineages were more closely related to each other than to the LF lineage (Fig. 4). More surprisingly, the results suggested that the three parental lineages were not monophyletic and cannot all be considered as extinct lineages. Indeed, the LF parental lineage and two extant outgroup species (*Sisymbrium irio* and *Orychophragmus violaceus*) seem to share a common ancestor, whereas the other two parental lineages (MF1 and MF2) belong to another monophyletic group, sister group to the former (Fig. 4). Such results can only be established when all subgenomes gene copies are present on a phylogenetic tree. Moreover, this result could explain the difficulties encountered in some studies to recover the Brassiceae monophyly with nuclear markers when Orychophragmus and species from the Sisymbridae clade were included in the dataset. These two lineages were indeed recovered nested within the Brassiceae (Warwick and Sauder 2005, Couvreur et al. 2010)

We found that the phylogenetic positions of the two outgroup species, although supported with high confidence, was not consistent among the trees obtained with alternative subgenomes annotation. Indeed, when using the annotation of Murat et al. (2015) or the concordant triplets dataset, the LF, MF1 and MF2 lineages appeared as a monophyletic group, placing the two extant outgroups as external lineages (Fig. 5). These strongly supported yet non-concordant results may be attributable to incomplete lineage sorting along the short internal branches of the trees, so that the result could differ across subsets of analyzed genes. Phylogenetic concordant methods could help assessing the discordance level between gene trees and estimates the proportion of genes supporting one or the other hypothesis (e.g. Ané et al. 2007). The LF genome could as well result from an ancient homoploid hybridization between the ancestors of the LF lineage and of the MF1/MF2 lineage. This homoploid hybridization preceding the allopolyploid event is another phenomenon that could generate discrepancies in the phylogenetic position of the parental subgenomes LF with respect to the outgroups *O. violaceus* and *S. irio*, depending on the parental origin of the gene sequenced. As illustrated in some other studies (e.g. Glemin et al. 2019) complex evolutionary histories of lineages such as incomplete lineage sorting or homoploid hybridization can blur the phylogenetic signal even if genome annotations are correct.

### A Fully Resolved Nuclear Phylogeny of Brassiceae Subtribes

Our results represent the first phylogenetic study of the mesohexaploid Brassiceae tribe that uses a large number of nuclear gene sequences and representatives from all major subtribes. In our results, the Brassiceae representatives appear as a monophyletic group (BP=100, PP=1) in all inferred phylogenies, which confirms previous results (Warwick and Black 1997; Hall et al. 2011; Arias and Pires 2012; Willis et al. 2014). We also confirm that all species of the tribe share the mesohexaploid event first characterized in the *Brassica rapa* genome (Wang et al. 2011a), as suggested by Lysak et al. (2007). Bootstrap values and posterior probabilities both strongly support the relationships among the major clades of the Brassiceae tribe (Figs. 2 and 4). The first diverging lineages of the tribe are, successively, Vella, Zilla and then Savignya. The basal position of the three clades is congruent with earlier studies based on chloroplast markers (Warwick and Black 1994; Warwick and Sauder 2005; Hall et al. 2011; Willis et al. 2014). However, the present results are different from those obtained with a drastically lower number of nuclear markers that systematically reached a different topology, possibly as a consequence of some confusion between homoeologs and proper orthologs (Warwick and Sauder 2005; Hall et al. 2011; Willis et al. 2014). Together with those of Willis et al. (2014), our results do not support the close relationship of the Savignya clade with the Oleracea clade, proposed by Arias and Pires (2012). Our results also clarify the relationship between the Nigra and Oleracea clades, which appear as sister clades, with Crambe as the first external lineage and then Cakile. Finally, in our results, the Raphanus genus (radish) belongs to the Nigra clade instead of the Oleracea clade as reported in many studies (e.g. Willis et al. 2014). As discussed by Yang et al. (2002), all phylogenies obtained with chloroplast data tend to place Raphanus within the Oleracea clade, whereas those based on nuclear genes place the genus within the Nigra clade (as confirmed here) (Yang et al. 1998; Yang et al. 1999; Hall et al. 2011), suggesting that the genus evolved after an homoploid hybridization between members of both clades, with the mother arising from the Oleracea clade.

Overall, our results demonstrate that taking into account the mesopolyploid origin of a clade (here, the Bassiceae tribe) is of central importance for proper phylogenetic analysis, altering some basic conclusions on the evolutionary history of the clade considered. The framework we have developed in the present study avoids conflating homoeologs and orthologs and can be applied to any group of species in which subgenome annotation is available. We believe that it will contribute to improve our understanding of the origins of polyploid lineages.

## Supplementary Material

Data available from the Dryad Digital Repository: http://dx.doi.org/10.5061/dryad.[NNN] and TreeBASE https://treebase.org/.

## Acknowledgements

We thank Hélène Martin for help with bioinformatics analyses, Nina Hautekeete and Yves Piquot for their contribution to plant sampling and Sara L. Martin from the Eastern Cereal and Oilseed Research Centre (Canada) who supplied Brassiceae seeds.

