## Supplemental material for "A New Tree-Based Methodological Framework to Infer the Evolutionary History of Mesopolyploid Lineages: An Application to the Brassiceae Tribe (Brassicaceae)"

**Online Appendix 1.** List of species used for RNA sequencing using Illumina Hiseq technology and information on sequencing. The number of paired-end reads was assessed using FastQC. The range for each species (min-max) is reported.

| Species | Source of seeds[a] | Organs (Number of individuals) | Illumina HiSeq technology | Read size | Number of raw reads | Number of clean reads |
|---|---|---|---|---|---|---|
| *Orychophragmus violaceus* | BGBe | Flower buds (2) | 2000, paired-end | 100bp | 14,151,140 - 14,976,841 | 12,920,785 - 13,756,251 |
| *Carrichtera annua* | LBG<br>BCN 3548 | Flower buds (3) | 3000, paired-end | 150bp | 19,186,051 - 34,288,654 | 17,927,018 - 32,416,939 |
| *Schouwia purpurea* | GCC no. 5780-81<br>BCN 8087 | Flower buds (3) | 3000, paired-end | 150bp | 36,054,912 - 39,167,535 | 34,141,935 - 36,712,467 |
| *Zilla spinosa subsp macroptera* | GCC no. 0731-67<br>BCN 8055 | Leaves (2) | 3000, paired-end | 150bp | 7,070,061 - 8,074,587 | 6,648,568 - 7,598,227 |
| | | Flower buds (1) | 3000, paired-end | 150bp | 43,184,474 | 40,639,896 |
| *Psychine stylosa* | BGB no.133<br>BCN 3515 | Flower buds (2) | 3000, paired-end | 150bp | 28,874,630 - 39,162,643 | 27,076,377 - 36,886,118 |
| *Crambe maritima* | Embouchure de la Slack, Ambleteuse (Pas-de-Calais) | Flower buds (3) | 3000, paired-end | 150bp | 22,250,949 - 30,817,974 | 20,746,799 - 28,941,942 |
| *Cakile maritima* | Digue du Braek, Grande-Synthe (Nord) | Flower buds (12) | 2500, paired-end | 100bp | 10,829,811 - 21,637,768 | 10,207,438 - 20,097,777 |

[a]BGBe, National Botanic Garden of Belgium, Meise, Belgium; GCC, Gómez-Campo Collection, Instituto Nacional de Investigaciones Agrarias, Madrid, Spain ; LBG, Leipzig Botanical Garden, Leipzig, Germany; BGB, Botanic Garden, Barcelona, Spain. BCN indicates the collection number on herbarium specimens deposited at Herbarium, Agriculture and Agri-Food Canada, Ottawa, Ontario, Canada (DAO).

**Online Appendix 2.** Summary statistics for transcriptome assemblies (after CAP3). All statistics were assessed using QUAST and are based on TRINITY contigs of size ≥ 500bp, unless otherwise noted. The range for each species (min-max) is reported, except for *Zilla spinosa subsp macroptera* for which the first line corresponds to the range of the two leaves samples and the second line corresponds to values of the single sample of flower buds.

| Species | Total number of TRINITY contigs (≥0bp) | Total number of TRINITY contigs | Total length (bp) | Largest contig (bp) | N50 (bp) | GC (%) |
|---|---|---|---|---|---|---|
| *Orychophragmus violaceus* | 65,359 - 78,128 | 39,529 - 48,912 | 61,313,006 – 75,284,383 | 11,290 - 14,979 | 1,821 - 1,857 | 42.56 - 42.65 |
| *Carrichtera annua* | 70,141 - 77,478 | 40,135 - 43,471 | 57,324,639 - 63,184,578 | 14,393 - 15,537 | 1,699 - 1,757 | 42.11 - 42.27 |
| *Schouwia purpurea* | 89,921 - 115,887 | 48,474 - 57,725 | 70,679,064 - 79,652,025 | 15,483 - 15,546 | 1,658 - 1,754 | 42.57 - 42.85 |
| *Zilla spinosa subsp* | 52,564 - 56,589 | 28,710 - 30,767 | 38,803,080 - 41,875,300 | 13,332 - 14,382 | 1,596 - 1,606 | 43.29 - 43.47 |
| *macroptera* | 103,503 | 55,420 | 80,126,538 | 15,556 | 1,745 | 42.30 |
| *Psychine stylosa* | 62,623 - 67,639 | 34,306 - 37,900 | 49,067,347 - 55,056,638 | 15,596 - 15,778 | 1,697 - 1,723 | 43.14 - 43.40 |
| *Crambe maritima* | 85,592 - 114,672 | 46,789 - 59,193 | 64,728,896 - 78,745,514 | 14,376 - 15,567 | 1,576 - 1,646 | 42.88 - 43.51 |
| *Cakile maritima* | 55,157 - 68,295 | 32,800 - 39,718 | 46,528,647 - 59,464,324 | 11,901 - 16,144 | 1,665 - 1,820 | 43.33 - 43.84 |

**Online Appendix 3.** Percentage of alignment gaps for each species in the concatenated alignments (ALL, LF, MF1 and MF2) constructed using the Liu database, calculated as the total number of alignment gaps relative to the total alignment length (bp). For each species, the reported values mirror together sites with missing information and indels, except for *A. thaliana* where the reported values mirror only the indels as we assume that there is no missing information in the TAIR database.

| | | Alignment gaps (%) | | | |
|---|---|---|---|---|---|
| Species | Data type | ALL | LF | MF1 | MF2 |
| *Arabidopsis thaliana* | genomic cds | 0.80 | 0.78 | 0.82 | 0.78 |
| *Brassica nigra* | genomic cds | 19.33 | 19.14 | 18.67 | 20.19 |
| *Brassica oleracea* | genomic cds | 9.31 | 9.36 | 9.00 | 9.58 |
| *Brassica rapa* | genomic cds | 4.83 | 3.56 | 5.56 | 5.40 |
| *Raphanus sativus* | genomic cds | 29.88 | 24.77 | 30.74 | 34.31 |
| *Raphanus raphanistrum* | genomic cds | 43.64 | 40.21 | 47.70 | 43.08 |
| *Eutrema salsugineum* | genomic cds | 10.86 | 10.62 | 10.92 | 11.06 |
| *Schrenkiella parvula* | genomic cds | 12.79 | 12.84 | 12.82 | 12.72 |
| *Sisymbrium irio* | genomic cds | 23.34 | 23.38 | 23.03 | 23.61 |
| *Orychophragmus violaceus* | transcriptome assembly | 20.41 | 20.58 | 20.19 | 20.46 |
| *Cakile maritima* | transcriptome assembly | 48.34 | 37.99 | 54.23 | 53.10 |
| *Crambe maritima* | transcriptome assembly | 53.78 | 48.32 | 54.87 | 58.34 |
| *Carrichtera annua* | transcriptome assembly | 65.73 | 57.28 | 70.82 | 69.33 |
| *Pychine stylosa* | transcriptome assembly | 61.02 | 51.24 | 64.31 | 67.82 |
| *Schouwia purpurea* | transcriptome assembly | 56.12 | 46.38 | 59.13 | 63.17 |
| *Zilla macroptera* | transcriptome assembly | 55.14 | 44.85 | 58.83 | 62.07 |

**Online Appendix 4.** Number of LF, MF1 and MF2 gene sequences collected in each investigated Brassiceae species in the final set of the 867 homolog alignments used for phylogenetic reconstruction.

| Species | Number of gene sequences in each subgenome | | |
|---|---|---|---|
| | **LF** | **MF1** | **MF2** |
| *B. rapa* | 851 | 840 | 839 |
| *B. oleracea* | 817 | 816 | 806 |
| *B. nigra* | 722 | 718 | 697 |
| *R. sativus* | 672 | 623 | 591 |
| *R. raphanistrum* | 553 | 493 | 523 |
| *C. maritima* | 549 | 428 | 433 |
| *S. purpurea* | 492 | 381 | 339 |
| *Crambe maritima* | 470 | 421 | 386 |
| *P. stylosa* | 432 | 326 | 299 |
| *Z. spinosa subsp macroptera* | 501 | 384 | 346 |
| *C. annua* | 376 | 265 | 269 |

**Online Appendix 5.** *Influence of the Subgenome Annotation Database on Phylogenetic Inferences.*

The 472 gene triplets from *Brassica rapa* reported by Murat et al. (2015) were obtained as well as 420 corresponding orthologous sequences from *Arabidopsis thaliana*. After filtering out the sequences shorter than 500bp, we obtained 325 triplets. 2 triplets were discarded after phylogenetic filtering on homolog trees constructed from the alignments of the three sequences of *B. rapa* and the sequence of *A. thaliana* (see Methods), giving a dataset of 323 homolog datasets. For each of the 323 homolog datasets, the three copies of *B. rapa* were mapped onto (i) the transcriptome assemblies and (ii) the genomic coding sequences of *Raphanus sativus*, *R. raphanistrum* and *Brassica nigra*, and the single copy of *A. thaliana* was mapped onto the genomic coding sequences of the outgroup species, in order to obtain full homolog datasets (Fig. 1a). After alignment of each of the 323 homolog datasets, homolog trees were constructed and 64 were discarded according to phylogenetic filters (see Methods). Orthologs' subtrees were identified in each of the 259 remaining homolog trees (Fig. 1b). On the 259 homolog alignments, 63 were discarded because of the percentage of Brassiceae sequences that were not included in an orthologs' subtree (196 remaining homolog alignments). For the analysis using the 181 'concordant triplets', 26 homolog groups were filtered out (length and phylogenetic filters) and 35 were discarded due to the percentage of Brassiceae sequences that were not included in an orthologs' subtree (120 remaining homolog groups). LF, MF1, MF2, "concatenated" and "homologous" matrices as well as species trees were constructed following the same procedure as for the Liu database (Liu et al. 2014) (Fig. 1c to 1e). Concerning the Bayesian inferences, for each of the five alignments obtained with both datasets (Murat and CT databases), two runs of 75,000,000 generations were completed with four chains each and trees were sampled every 2000 generations.

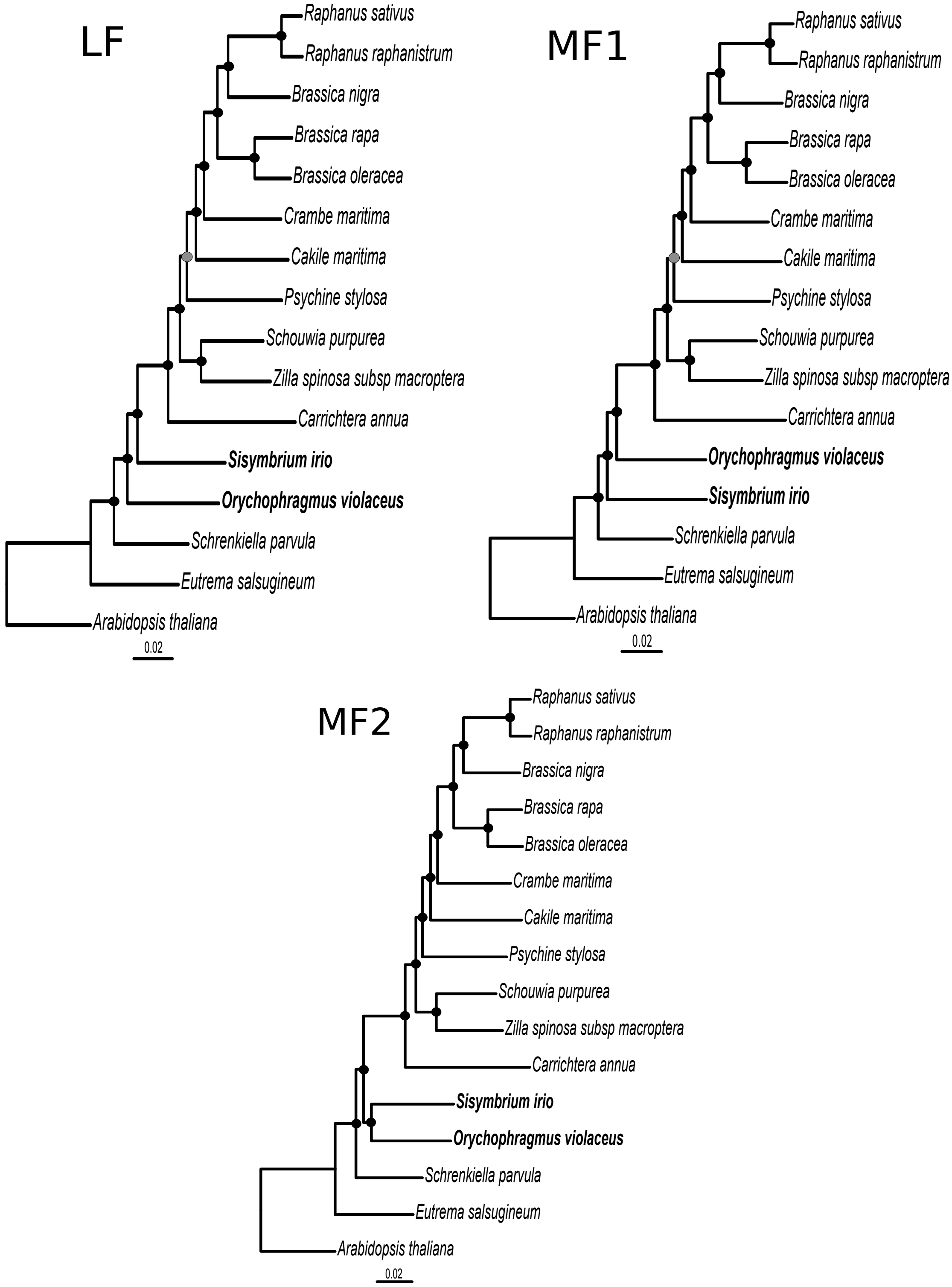


**Online Appendix 6.** Maximum likelihood phylogeny of the tribe Brassiceae and its relatives from the analyses of the LF, MF1 and MF2 alignments based on **Liu et al (2014) genes annotation**. Black circles indicate nodes with maximal support (BP = 100 / PP = 1.0). Smaller support values are indicated in grey (90 ≤ BP <100 and PP=1). Brassiceae' closest outgroups are in bold.

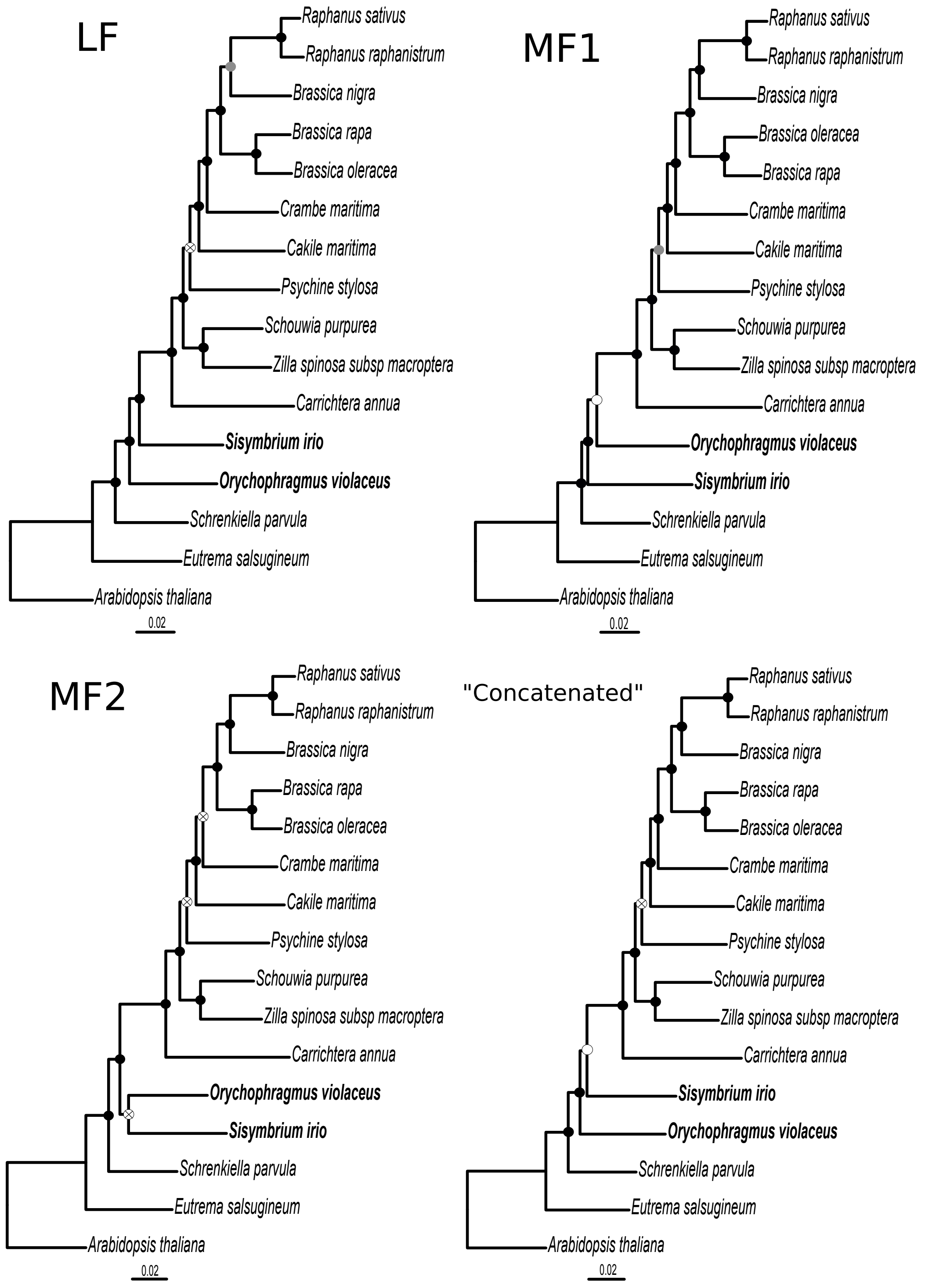


**Online Appendix 7.** Maximum likelihood phylogeny of the tribe Brassiceae and its relatives from the analyses of the LF, MF1, MF2 and « concatenated » **filtered alignments**. Black circles indicate nodes with maximal support (BP = 100 / PP = 1.0). Smaller support values are indicated with grey (90 ≤ BP <100 and PP=1) and white (80 ≤ BP < 90 and PP=1) circles. Supports with a BP < 80 and a PP=1 are indicated with a cross. Brassiceae' closest outgroups are in bold.

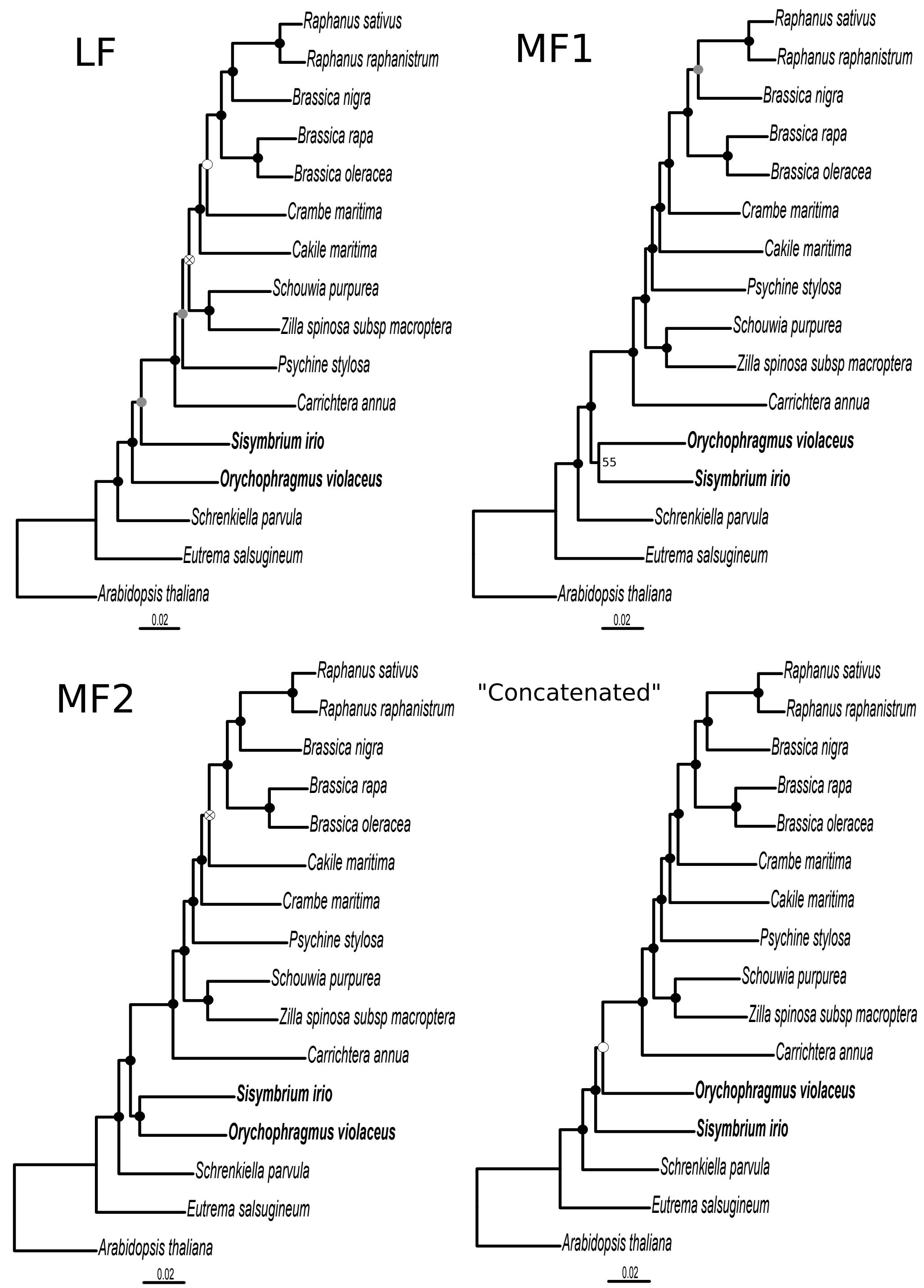


**Online Appendix 8.** Maximum likelihood phylogeny of the tribe Brassiceae and its relatives from the analyses of the LF, MF1, MF2 and "concatenated" alignments based on **Murat et al (2015) genes annotation**. Black circles indicate nodes with maximal support (BP = 100 / PP = 1.0). Smaller support values are indicated with grey (90 ≤ BP <100 and PP=1) and white (80 ≤ BP < 90 and PP=1) circles. Supports with a BP < 80 and a PP=1 are indicated with a cross. For the MF1 reconstruction there was an incongruence between the ML and the Bayesian trees: either *S. irio* and *O. violaceus* were monophyletic (BP= 55) or *O. violaceus* was closer to the Brassiceae (PP=1). Brassiceae' closest outgroups are in bold.

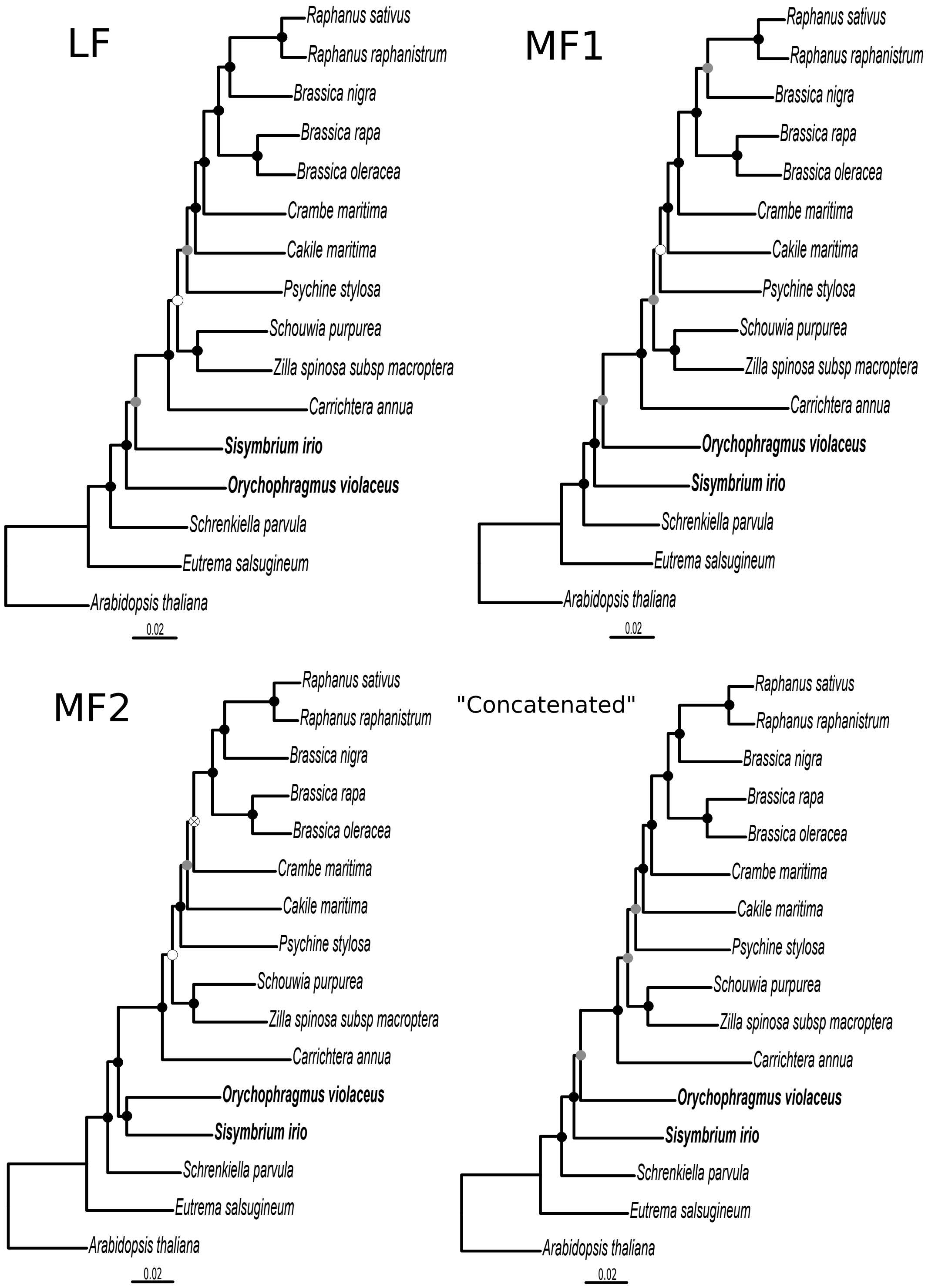


**Online Appendix 9.** Maximum likelihood phylogeny of the tribe Brassiceae and its relatives from the analyses of the LF, MF1, MF2 and “concatenated” alignments from the **“concordant triplets” genes annotation**. Black circles indicate nodes with maximal support (BP = 100 / PP = 1.0). Smaller support values are indicated with grey (90 ≤ BP <100 and PP=1) and white (80 ≤ BP < 90 and PP=1) circles. Supports with a BP < 80 and a PP < 0.95 are indicated with a cross. Brassiceae’ closest outgroups are in bold.

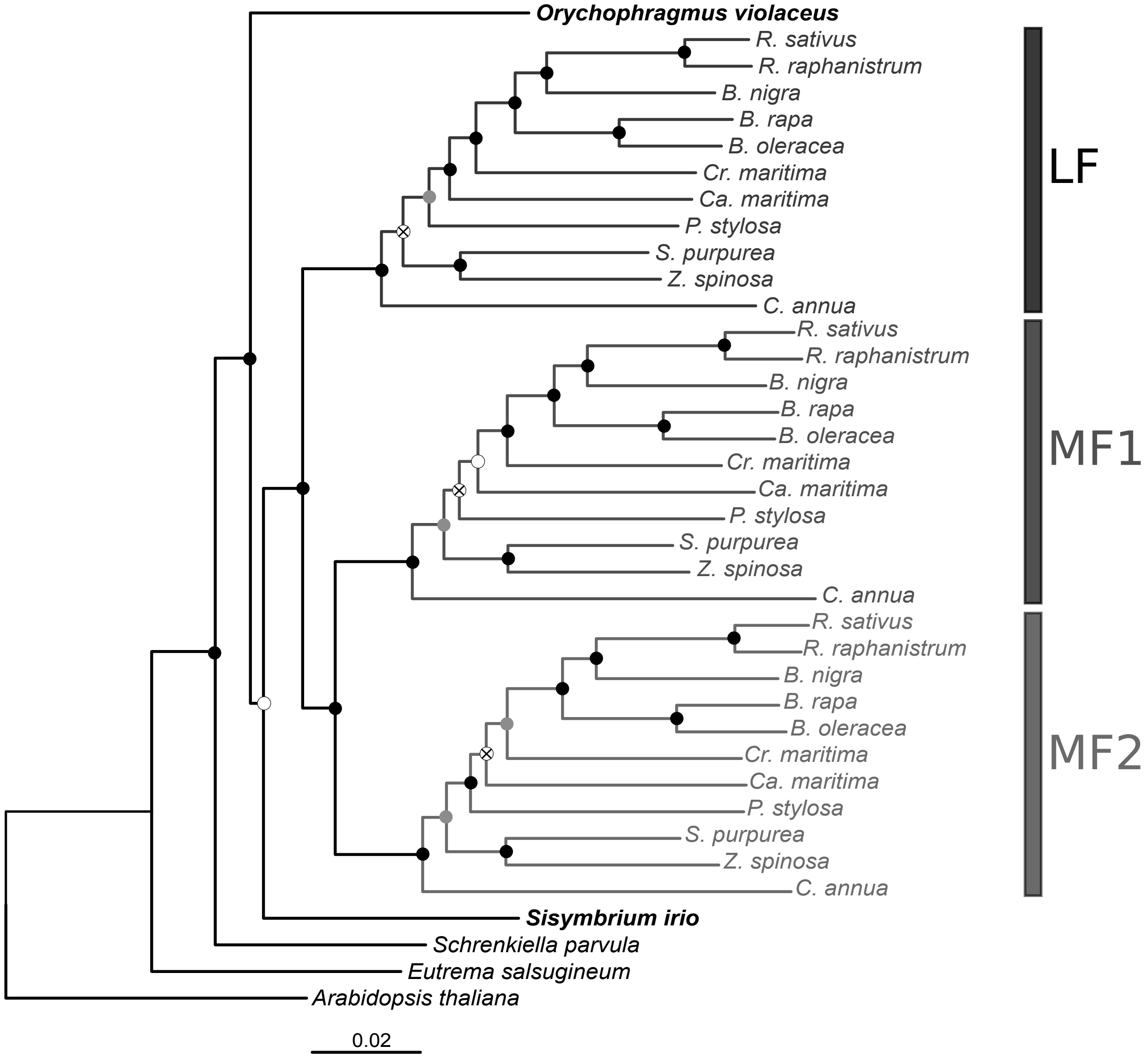


**Online Appendix 10.** Maximum likelihood phylogeny of the tribe Brassiceae based on the "homologous" alignment obtained with the Brassica **"concordant triplets" genes annotation**. The phylogeny displays the three sub-trees corresponding to the three subgenomes present in the Brassiceae species. Brassiceae' closest outgroups are in bold. Black circles indicate nodes with maximal support (BP = 100 / PP = 1.0). Smaller support values are indicated with grey (90 ≤ BP <100 and PP=1) and white (80 ≤ BP < 90 and PP=1) circles. Supports with a BP < 80 and a PP=1 are indicated with a cross.
